# Automated classification of estrous stage in rodents using deep learning

**DOI:** 10.1101/2022.03.09.483678

**Authors:** Nora S. Wolcott, Kevin K. Sit, Gianna Raimondi, Travis Hodges, Rebecca M. Shansky, Liisa A. M. Galea, Linnaea E. Ostroff, Michael J. Goard

## Abstract

The rodent estrous cycle modulates a range of biological functions, from gene expression to behavior. The cycle is typically divided into four stages, each characterized by distinct hormone concentration profiles. Given the difficulty of repeatedly sampling plasma steroid hormones from rodents, the primary method for classifying estrous stage is by identifying vaginal epithelial cell types. However, manual classification of epithelial cell samples is time-intensive and variable, even amongst expert investigators. Here, we use a deep learning approach to achieve classification accuracy at expert levels in a matter of seconds. Due to the heterogeneity and breadth of our input dataset, our deep learning approach (“EstrousNet”) is highly generalizable across rodent species, stains, and subjects. The EstrousNet algorithm exploits the temporal dimension of the hormonal cycle by fitting classifications to an archetypal estrous cycle, highlighting possible misclassifications and flagging anestrus phases (e.g., pseudopregnancy). EstrousNet allows for rapid estrous cycle staging, improving the ability of investigators to consider endocrine state in their rodent studies.

## INTRODUCTION

With the broad incorporation of female animals into previously all-male studies^1,2^ we are at a critical juncture for the interpretation of endocrine physiology. In naturally cycling humans, the menstrual cycle lasts 28 days and is characterized by defined peaks in steroid hormones such as estradiol and progesterone^3–9^. In female rats and mice, the analogous estrous cycle lasts 4-5 days^10^, but exhibits steroid hormone fluctuations similar to the 28 day human menstrual cycle. The estrous cycle was first described over a century ago^11^, yet the criteria for tracking this cycle remain subjective and variable between experimenters^12^. Determining the stage of estrous is critical to evaluating the state of the hypothalamic-pituitary-ovarian axis, which has implications in a myriad of factors including gene expression^13,14^, neuronal structure and connectivity^3,15^, and pharmacological efficacy^16^. In addition, correct interpretation of estrous stage is useful for timed pregnancy in rodents and changes in cycle regularity can be used as a proxy for changes in other critical hormones such as corticosterone^17,18^.

The estrous cycle can be divided into four stages: diestrus, proestrus, estrus, and metestrus^19–23^. While techniques such as vaginal opening evaluation, vaginal wall impedance, and urine biochemistry have all been used as methods for determining estrous stage^20^, epithelial cell cytology remains the most common and reliable strategy^6,9,10,13^. Classification using vaginal cytology is typically performed by manually counting or estimating the relative prevalence of epithelial cell types, including leukocytes, cornified epithelial, and nucleated epithelial cells, and using the proportionality of these subtypes to determine stage^10,19^.

Despite the prevalence of this method, there are several limitations of epithelial cell cytology for estrous stage classification: 1) it is time consuming and requires extensive training. 2) it lacks generalizability; even expert classifiers may have trouble generalizing across rodent species, stains, and subjects. 3) it is inconsistent between labs, as classification can vary widely between human examiners^12^. Here, we address these challenges using a novel deep learning algorithm that can generate estrous stage classifications on the order of seconds.

Convolutional neural networks (CNNs) have outperformed human experts in diagnosing retinal disease^24^, skin cancer^25^, syndromic genetic diseases^26^, and a host of other medical conditions^27^. These networks are broadly useful for their speed and reliability. Although CNNs are difficult to train from scratch, requiring massive training data sets for accurate classification, transfer learning can exploit the multilayered architectures of pretrained networks to classify complex biological images^28,29^.

Here, we have compiled a large-scale multi-laboratory dataset of cytology images (“EstrousBank”). We then used EstrousBank to train a deep learning algorithm (“EstrousNet”) to effectively recognize structural markers of the estrous cycle in a manner generalizable across subjects, stains, and rodent species. The resulting classifications are not significantly different than expert human examiners in any stage surveyed. The predictions generated by EstrousNet can be enhanced by using sequentially collected data to fit cytological samples with an archetypal estrous cycle. Cycle fitting, along with training, classification, and output, are operated through an interactive graphical user interface (GUI). Taken together, these results show that our deep learning approach is capable of rapid and accurate classification of estrous stage.

## RESULTS

### EstrousBank: an open resource for analysis of vaginal cytology images

A major barrier to the development of software to analyze the estrous cycle is a data-poor environment that requires experimenters to collect their own cytology images. In our efforts to make the EstrousNet algorithm generalizable across groups, we have compiled the largest known image bank of estrous cytology images. EstrousBank currently spans five labs, five stains, two magnifications, and multiple rodent species (**Fig. 1A-C, Supplementary Table S1**). The complete image bank comprises 12,719 vaginal cytology images and is freely available for analysis by outside laboratories. We will continue to add to the image bank as more samples become available. Cytological samples across labs were collected using a standard lavage or swabbing procedure (See Methods). Briefly, epithelial cells were exfoliated from the superficial vaginal cavity via sterile saline and transferred to a glass microscope slide. Samples were allowed to dry for up to 24 h before staining with one of several compounds, and images were collected using brightfield microscopy at a range of magnifications (**Supplementary Table S1**).

**Figure 1.**
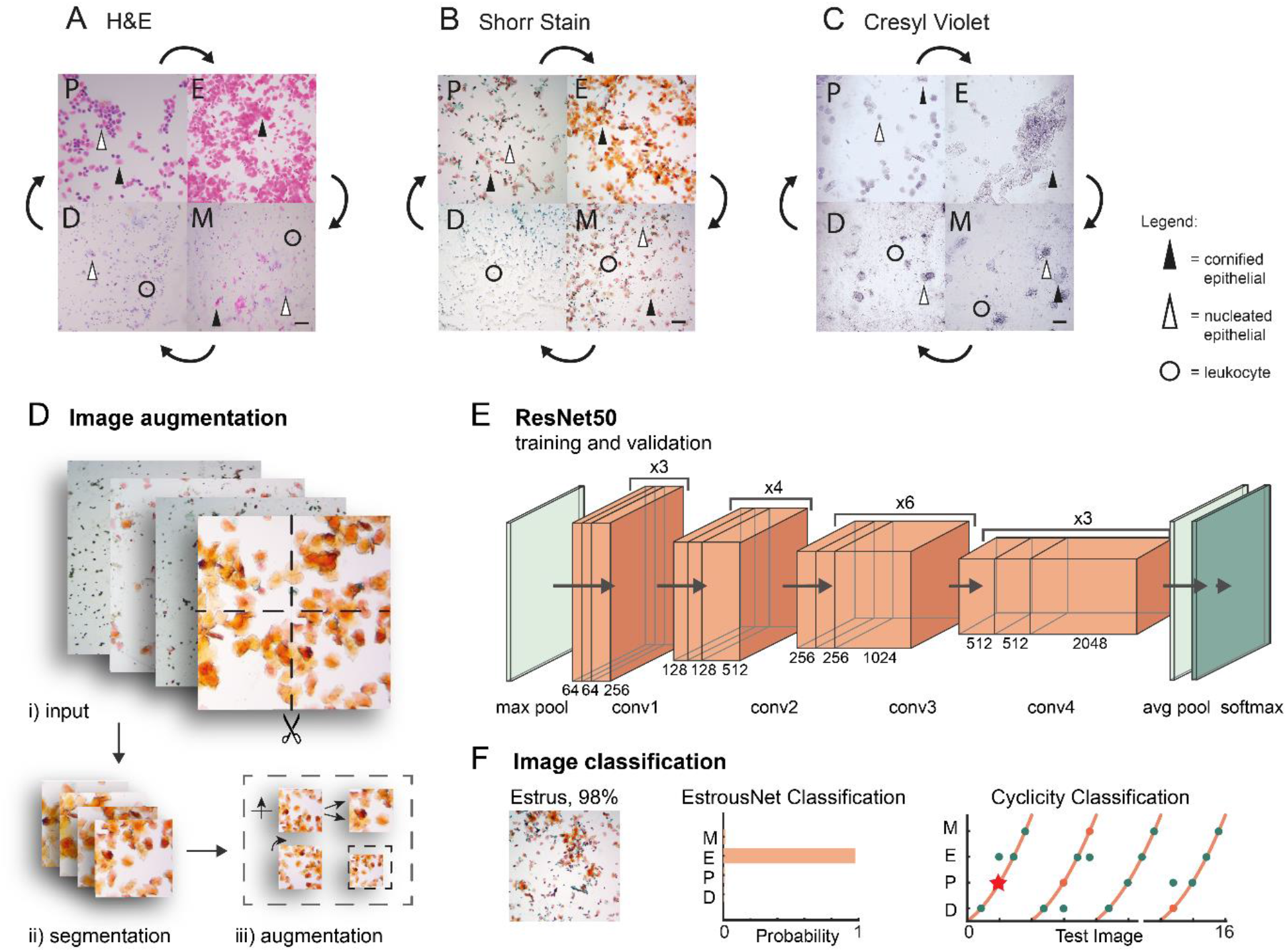
General schematic of the EstrousNet pipeline and representative cytological images. **A**. Hematoxylin and eosin (H&E)-stained vaginal cytology from a wild-type Sprague Dawley rat (Ostroff lab). Scale bar = 10 μm. **B**. Shorr-stained vaginal cytology from a Slc7a7-cre x TIT2L-GCaMP6s Bl6 mouse (Goard lab). Scale bar = 10 μm. **C**. Cresyl violet-stained vaginal cytology from a wild-type Long-Evans rat (Galea lab). Scale bar = 10 μm. **D**. Image augmentation schematic: images are first quadrisected, then reflected, scaled, rotated, and translated in our preprocessing pipeline. **E**. The base architecture of ResNet50 that is used for the transfer learning algorithm. Processed input images are transferred to a max pooling layer. Then, the images are processed through four convolutional units, which converge onto custom pooling and SoftMax classification output layers. **F**. Schematic of the EstrousNet GUI output. Estrous stage classifications are generated from the deep learning network, and the cycle tracking algorithm flags potential outliers.

EstrousBank contains images from all four stages of the estrous cycle, which were classified by experts according to classical cytology parameters, which are as follows^20–23^: mouse diestrus is characterized by an abundance of small leukocytes, a sharp decrease in proportions of keratinized anucleated epithelial cells, and lower numbers of both small and large nucleated epithelial cells (**Fig. 1A-C**). Mucosal secretions appear thick and stringy when present. Proestrus is a more transient stage characterized by a uniform spread of small rounded basophilic nucleated epithelial cells, and low proportions of anucleated cornified epithelial cells (**Fig. 1A-C**). Estrus is typically identified by the high proportion of large anucleated cornified epithelial cells, which often form clumps or sheets that become more prominent in late estrus (**Fig. 1A-C**). Metestrus is a short stage identified by the presence of both nucleated epithelial and cornified epithelial cells, with leukocytes clustered around them, and an elevated level of mucosal secretions (**Fig. 1A-C**). While others have broken down diestrus into 2-3 substages, here we consider metestrus to be its own distinct stage preceding diestrus. These characterizations are largely consistent between mice and rats, but the following differences have been observed: rats exhibit a higher proportion of large ovular nucleated epithelial cells in late estrus, shorter periods of proestrus/metestrus, and lower proportions of anucleated cornified epithelial cells in metestrus^21^. Given these similarities, we trained EstrousNet on cytology images from several strains of mice and rats to improve generalization across model systems; with 34.1% of the image set from mice and 65.9% from rats.

Although previous studies have used computational methods to analyze vaginal cytology^12,30^, the input datasets for these networks have historically been restricted to a single stain. To further enhance generalizability, the training and validation image sets for EstrousNet include samples stained with H&E, Shorr, Giemsa, cresyl violet, and crystal violet stains, at magnifications of 10x and 20x (**Fig. 1A-C, Supplementary Table S1**). The resulting pretrained CNN is highly generalizable and effective in classifying low resolution images and those containing debris from vaginal swab.

### A ResNet50-based CNN architecture maximizes EstrousNet performance

To classify estrous stage from vaginal cytology images, we developed a classification pipeline using a convolutional deep learning network to detect cell boundaries and recognize endocrine biomarkers within cytological samples. For training and validation, we used consensus classifications (see Methods) to attach an estrous stage label to each image. EstrousNet is trained on subsets of EstrousBank images that are augmented for a greater volume of training data. Input images are first segmented into quadrants (**Fig. 1D.i, ii**), then reflected, rotated, scaled, and translated within the Net (**Fig. 1D.iii**). The augmented images undergo luminance normalization, then are converted to 3-channel grayscale arrays for more efficient feature extraction (**Supplementary Fig. S1**). Next, these augmented images are compiled into a large datastore and fed into the ResNet50 architecture, which consists of four convolutional stages of increasing dimension (**Fig. 1E**). The convolutional layers of the network converge on a SoftMax classification layer, which outputs probabilistic classification of estrous cycle stage (**Fig. 1E, F**). This classification is optionally supplemented by fitting the test images to a curve describing the length and phase of the estrous cycle (**Fig. 1F**). For images in which the cyclicity prediction and net prediction disagree, the interactive GUI will ask the user to select which classification to use. The composite classifications of the EstrousNet and cyclicity predictions provide the experimenter with an informed estrous stage classification.

Previous studies investigating the efficacy of transfer learning in biological tissue classification have used several CNN architectures^12,28,29^. Here, we evaluated four different pretrained networks: VGG-19, Inception v3, MobileNet V2, and ResNet-50 (**Fig. 2A**)^31–34^. Each base architecture was originally trained on more than one million images from the ImageNet database and retrained on an augmented dataset made up of 80% of EstrousBank images, with 10% of images reserved for validation and 10% reserved for testing (**Fig. 2B, C**). All base architectures have previously been used for supervised learning in biological classification tasks and achieved accuracy comparable to or exceeding that of human coders^24–27^. The mean validation accuracies averaged over 3 iterations for each architecture are as follows: VGG-19 = 79.7%, Inception v3 = 77.5%, MobileNetV2 = 65.5% and ResNet-50 = 88.9**%** (**Fig. 2A**). These accuracies are calculated based on ground truth data defined by benchmark classifications between 3 expert human examiners. Based on these results, we concluded that ResNet-50 was the most effective architecture.

**Figure 2.**
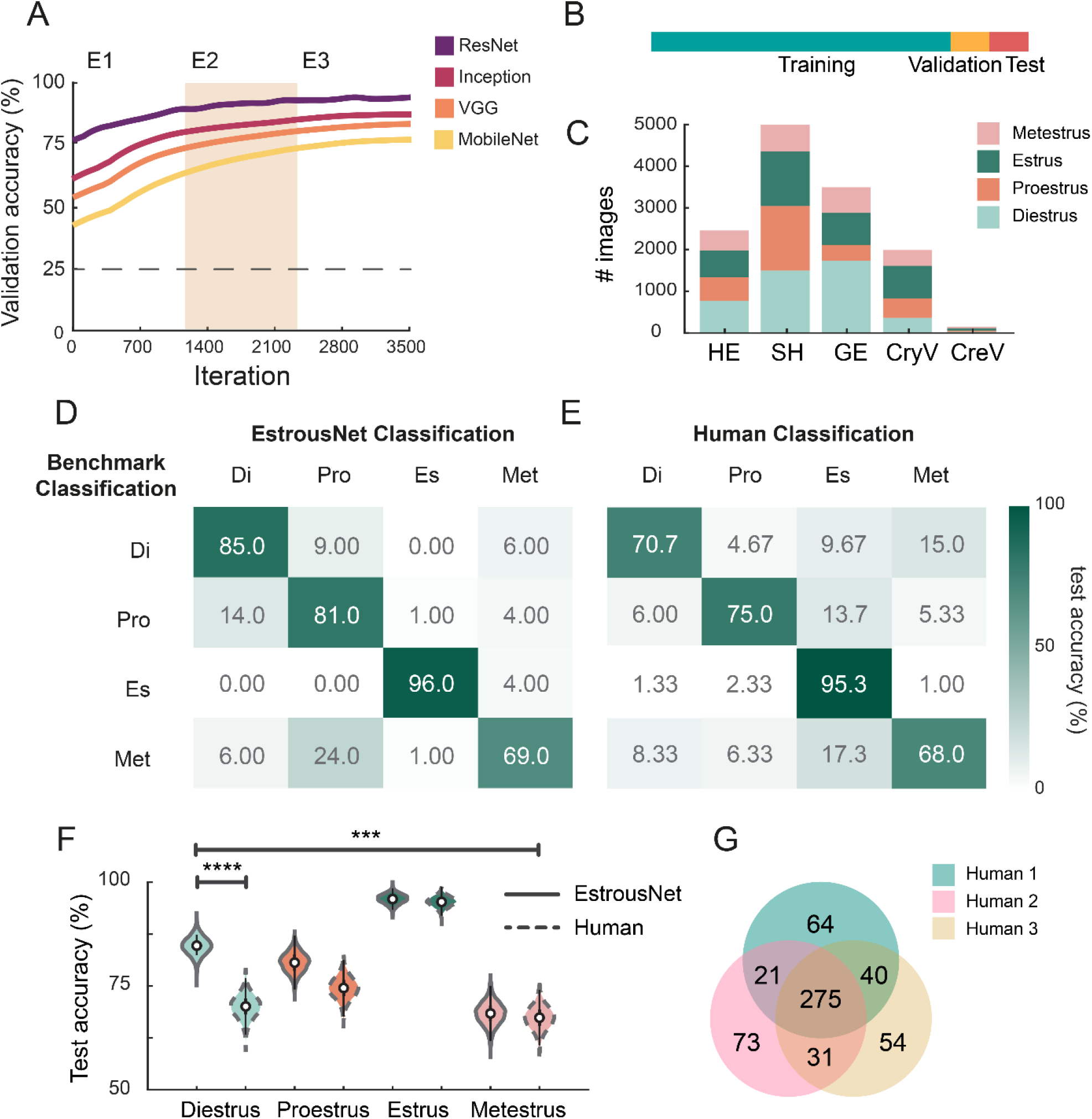
EstrousNet accuracy is comparable to human experts. **A**. Validation accuracy curves for EstrousNet trained using four different base architectures: ResNet50, Inception v3, VGG-19, and MobileNet v2. All networks were trained on EstrousBank images. Mean validation accuracy across 3 testing iterations. **B**. Schematic of the EstrousBank split for training, validation, and test sets. By percentage, this split is 80%, 10%, and 10%, respectively. **C**. Breakdown of EstrousBank by stain and stage. Stains from left to right are hematoxylin and eosin (HE), Shorr stain (SH), Giemsa stain (GE), crystal violet (CryV), and cresyl violet (CreV). The complete bank consists of *n* = 12,719 cytology images. **D**. Confusion matrix of EstrousNet classifications, represented here as a heatmap, with consensus from benchmark classification acting as our ground truth. Numbers represent the number of images classified for each stage, from a test set made up of 400 images (100 images from diestrus, proestrus, estrus, and metestrus). **E**. Confusion matrix of human classification, represented as a heatmap, with ground truth stages as described previously. **F**. Average test accuracy distributions in each estrous stage for EstrousNet vs human classifications. EstrousNet distributions are identified by a continuous line while human classifications are identified by a dotted line. Distributions were created by bootstrapping data over 50000 iterations, sampling without replacement. Error bars are 25th (75th) percentiles minus (plus) the interquartile range (75th percentile minus 25th percentile). Asterisks indicate significance as determined by Fisher’s Exact Test; diestrus: odds ratio = 0.68, 95% confidence interval = 0.55-0.83, *p* = 1.2 × 10^−5^, proestrus: odds ratio = 0.68, 95% confidence interval = 0.55-0.83, *p* = 0.075, estrus: *p* = odds ratio = 0.68, 95% confidence interval = 0.55-0.83, *p* = 0.84, metestrus: odds ratio = 0.68, 95% confidence interval = 0.55-0.83, *p* = 0.60. Across all stages accuracy was significantly different, with odds ratio = 0.68, 95% confidence interval = 0.55-0.83, *p* = 2.1 × 10^−4^, Fisher’s Exact Test. **G**. Venn diagram of the overlap between human expert coders, with a total of 400 classifications for each coder.

### EstrousNet outperforms human coders in both speed and accuracy

The cytology images in our training set were originally sorted into stages by expert human classifiers. These classifications were made using subjective assessments according to established approaches^10,20,21^ (see Methods). Unfortunately, human classification is limited by inter-experimenter variability and differences in experience with particular species, strains, and histological stains. In addition, the CNN may be capable of identifying subtle morphological features that are difficult for humans to identify, such as increased cell clumping in estrus and higher mucus content in metestrus and diestrus.

To quantify differences between EstrousNet and human coders, we compared classification performance on a test set of 400 randomly selected images (100 from each stage) between EstrousNet and three expert human coders. Across the test image set, EstrousNet classified stages significantly more accurately than human examiners (odds ratio = 0.68, 95% confidence interval = 0.55-0.83, *p* = 2.1 × 10^−4^; Fisher’s Exact Test). Breaking down performance by stage, EstrousNet achieved significantly greater accuracy than expert human examiners for diestrus (odds ratio = 0.6791, 95% confidence interval = 0.55-0.83, *p* = 1.2 × 10^−5^), whereas accuracy was higher, but not significantly different than expert examiners, for proestrus (odds ratio = 0.68, 95% confidence interval = 0.55-0.83, *p* = 0.075), estrus (odds ratio = 0.6791, 95% confidence interval = 0.55-0.83, *p* = 0.84) and metestrus (odds ratio = 0.68, 95% confidence interval = 0.55-0.83, *p* = 0.60; Fisher’s Exact Test for all comparisons; **Fig. 2D-F**). EstrousNet classifications also achieved impressive speed, with an average rate of 0.10 +/-0.005 s (mean +/-SE) per image.

Expert human staging showed a large degree of variance, with only 275 image classifications, or 68.75% of the total test set, shared between all three coders (**Fig. 2G**). A notable number of classifications, 15.9%, were unique to one human coder (**Fig. 2G**). Therefore, even amongst expert human classifiers, classifications can vary widely across a generalizable dataset of cytology images.

### EstrousNet is generalizable across species, stains, and subjects

To further quantify EstrousNet performance for each estrous stage, we measured the area under the receiver operating characteristic (auROC) for each stage independently. EstrousNet demonstrated auROC values greater than 0.79 for all four estrous stages, with estrus achieving the highest auROC at 0.98 (**Fig. 3A**). Despite this high performance, there are areas in which EstrousNet shows tendencies towards misclassification. Sensitivity and specificity curves show that EstrousNet is stronger in eliminating false negative results than false positive results, indicating a higher degree of sensitivity than specificity (**Fig. 3B**). For example, if EstrousNet is given an image of an unknown stage and asked if the sample is from an animal in diestrus, EstrousNet is more likely to classify the sample as diestrus when it is not (false positive), than to classify it as not diestrus when it is (false negative). Therefore, most misclassifications are specificity errors, which could potentially be reduced with further optimization.

**Figure 3.**
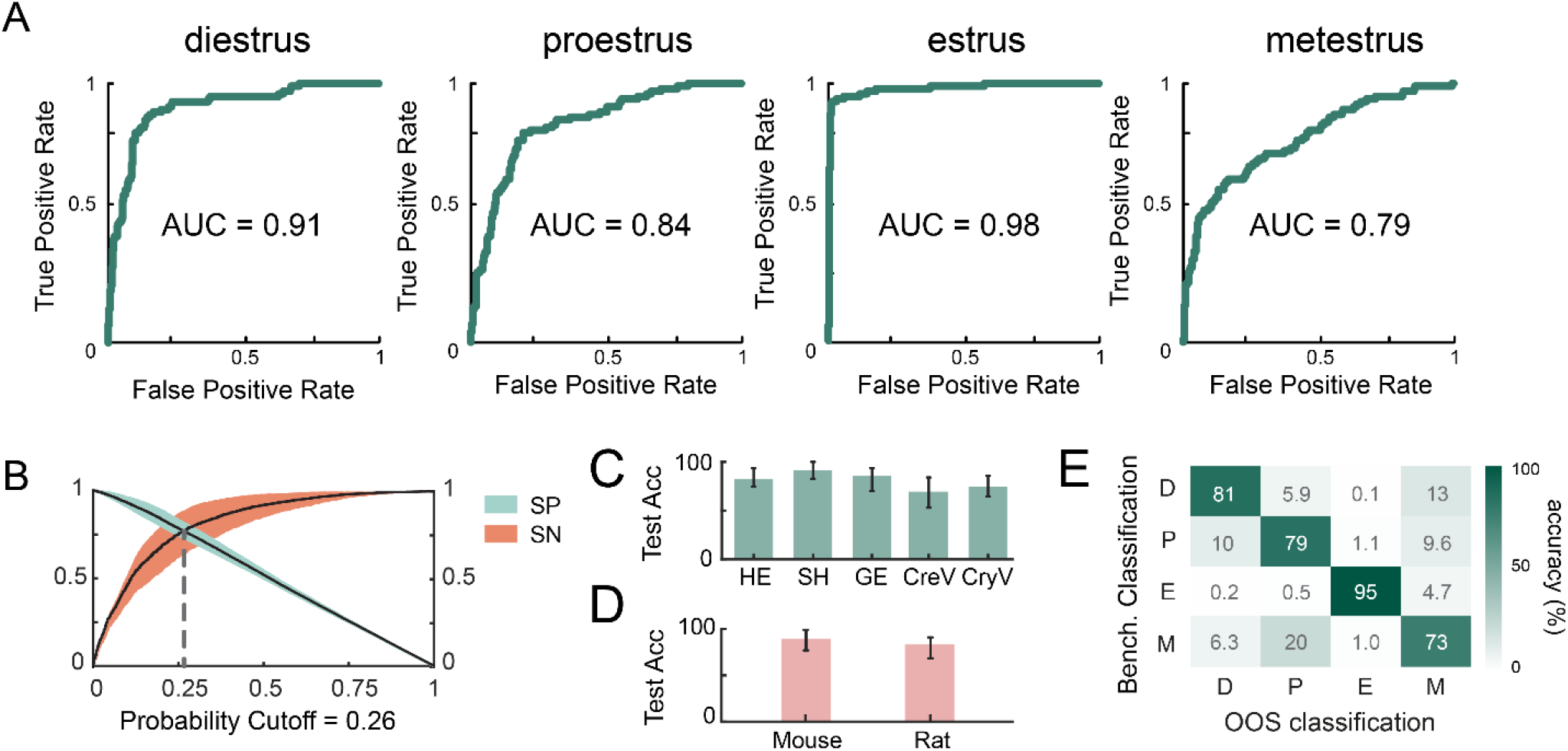
EstrousNet performs similarly across groups. **A**. auROC curves for each estrous stage. True positives for each stage are determined by benchmark classifications. **B**. Specificity (SP) vs sensitivity (SN) curves for EstrousNet, with the probability cutoff at 0.26 defined as the intersection between curves (dotted grey line). Standard error shown in orange and blue for sensitivity and specificity, respectively. **C**. Out of sample testing across 4 different stains: hematoxylin and eosin, Shorr stain, Giemsa stain, cresyl violet, and crystal violet. Test accuracy represented as a distribution across 1000 testing iterations, with mean % SE shown. Accuracy differences between stains are not significant (*F*(4,198) = 3.14, *p* = 0.10, one-way ANOVA). **D**. Out of sample testing between mouse and rat species. Test accuracy represented as a distribution across 1000 testing iterations, with mean % SE shown. Accuracy differences between species are not significant (*F*(1,198) = 7.87, *p* = 0.73, one-way ANOVA). **E**. Out of sample (OOS) classification for each stage of the estrous cycle between different animals, represented as a heatmap. Benchmark classification was used as a proxy for ground truth. K-fold cross-validation was used to estimate accuracy across stages, with k = 6 groups. Testing accuracy was averaged between each fold to generate the most unbiased estimate across all groups. Accuracy differences between subjects are not significant (*F*(5,198) = 6.98, *p* = 0.60, one-way ANOVA).

In out-of-sample trials in which the CNN was tested on different categories of unseen data, EstrousNet did not show significant differences in test accuracy between any of the given stains it was tested on, including H&E, Shorr, Giemsa, cresyl violet, or crystal violet (**Fig. 3C**). Additionally, despite cytological differences, images from mice and rats did not show significant differences in testing accuracy (**Fig. 3D**). Finally, cross-validation across 6 evenly split groups of subjects, including rats and mice of different strains, did not reveal any out-of-sample differences in test accuracy between animals (**Fig. 3E**).

### Using cycle fitting for predictive stage classification

When an experimenter classifies estrous stage from epithelial cytology, they not only consider cell morphology and relative prevalence, but also how images might correspond to a typical estrous cycle. Helpfully, some common confusion errors occur between stages that are temporally distinct. For instance, true metestrus is classified as proestrus at a rate of 24.0% despite being non-adjacent stages of the cycle (**Fig. 3D**). As a result, we can exploit the natural sequence of the estrous cycle to identify these errors when test images are taken consecutively. To this end, EstrousNet uses a predictive algorithm that fits an archetypal estrous cycle to the labels generated by the net and identifies outliers (**Fig. 4A, B**).

**Figure 4.**
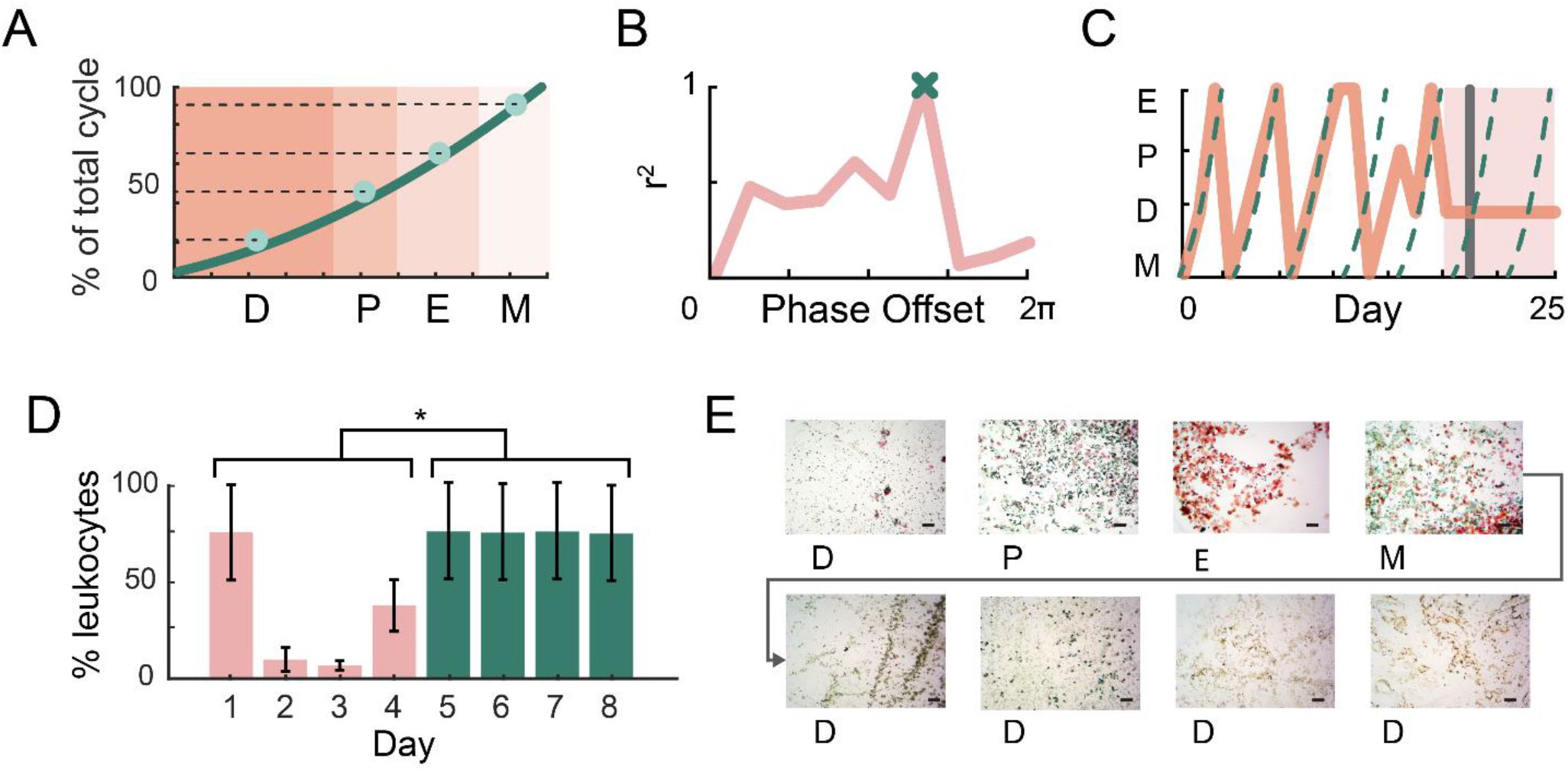
Sequential estrous classifications are fit to an archetypal cycle. **A**. Schematic of the custom waveform used for temporal cycle fitting. Color blocks indicate the length of each estrous stage as a percentile of average total cycle length, with a curve fitted to the midpoint of each stage (see Methods). Stage lengths are based on a consensus archetypal cycle from previous studies^10,18,20–23,30,35–38^. **B**. Pearson’s coefficient for each iterative fit of the custom waveform to an example 16-day cycle, at increments of 0.1 cycles. The best fit is determined by global maxima, marked by an ‘x’ for this example cycle. **C**. Example of a naturally cycling mouse tracked across 25 days, with the animal’s cycle shown as a solid orange line and the fitted cycle curve as a dotted teal line. The mouse initially exhibited regular cycles but entered pseudopregnancy on day 18 (shaded area), causing EstrousNet to give the user a pseudopregnancy warning message (grey line). **D**. Proportion of leukocytes in cytological cell counts before (blue) and after (pink) pseudopregnancy. Mean +/-SE, *F*(1,6) = 7.44, *p* = 0.034, as determined by two-way ANOVA. Asterisk indicates significance of *p* < 0.05. **E**. Cytology images from a normally cycling mouse entering pseudopregnancy, demonstrating prolonged diestrus, with an abnormally high proportion of leukocytes. Scale bars = 10 μm.

A custom cycle waveform was created based on the duration of estrous stages reported from thirteen groups^10,18,20–23,30,35–38^. If more than 4 days of test images are selected (i.e., n > 4*x where x is the sampling frequency per day), the algorithm can fit an archetypal cycle to the data to determine the relative phase that best fits the classification labels. The phase of this periodic waveform was shifted by increments of 0.1 cycles to find the best fit for the input data (**Fig. 4B**). We developed a MATLAB-based graphical user interface (GUI) that allows experimenters to select which stage to accept in cases where the net prediction and cyclicity predictions do not match (**Supplementary Fig. S2**).

Fitting stages to an archetypal cycle also allows us to identify disruptions in the estrous cycle, such as those observed when the rodent enters pseudopregnancy, a condition occasionally induced by vaginal swab or lavage^21,22^. Observations of anestrous stages are also useful for those inducing timed pseudopregnancy for reproductive management and embryo transfer^10,39^. To address this, EstrousNet will alert the user with a pseudopregnancy warning flag if the animal stays in diestrus for > 50% longer than in previous cycles (**Fig. 4C**). Manual cell counts from an example cycle in which a mouse was lavaged once a day for 8 consecutive days shows a significant increase in the proportion of leukocytes observed once the animal enters pseudopregnancy (**Fig. 4D**, *F*(1,6) = 7.44, *p* = 0.034). Such persistent diestrus following a cornified swab is consistent with previous observations of chemically or mechanically induced pseudopregnancy, and can be seen in a series of cytological images (**Fig. 4E**)^22^.

Additionally, cycle fitting may help to identify stages that do not fall into a traditional category. While here we refer to estrous as consisting of 4 substages, as many as 13 substages have been identified, each corresponding to physiologically distinct steroid hormone concentrations^41,42^. For the intermediate period(s) between each stage, manual cell counting of sequential samples revealed cell proportionalities distinct to these transition stages (**Fig. 5**). Despite these advancements, more sequential data will be needed for EstrousNet to reliably classify transition stages.

**Figure 5.**
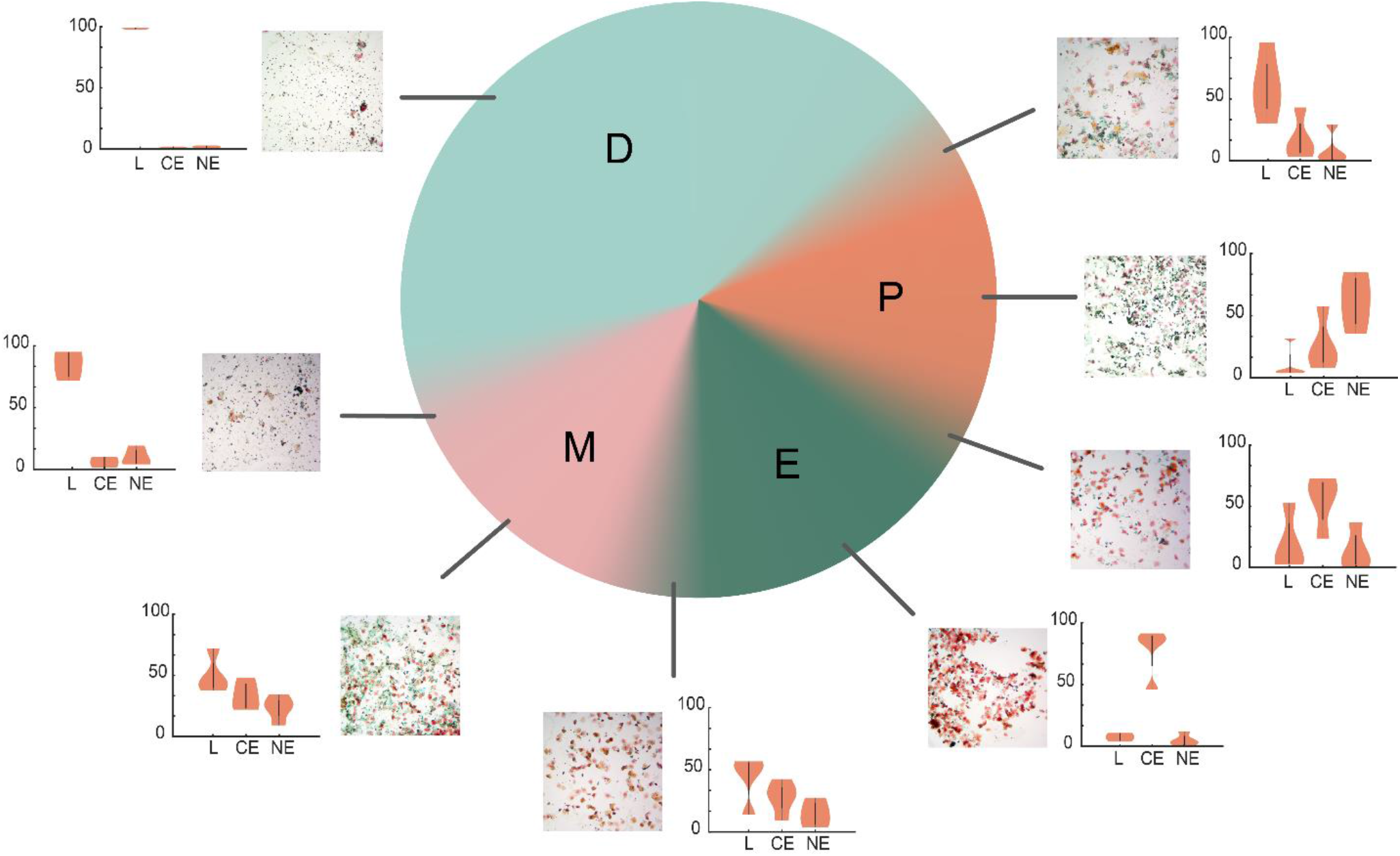
Characterization of cell types across the estrous cycle. A pie chart of the four estrous stages broken down by stage length across the 4–5-day cycle. Transition states between the four classical estrous stages are shown in gradient, and cytology collected from between the stages are included. Raw images and violin plots of cell counts from 8 primary and transition stages are shown (four examiners). The violin plots indicate the proportions of leukocytes, cornified epithelial, and nucleated epithelial cells, respectively (mean +/-SE).

## DISCUSSION

Here, we created a deep learning network for automated classification of estrous stage. The 12,719 images that constitute EstrousBank allow us to classify the four stages of estrous in a manner generalizable to stain, subject, and rodent species. EstrousBank is a valuable tool for future developers in the rapidly advancing machine learning field, and the benchmark classifications within the bank provide a guide for those learning to identify estrous stage. Our EstrousNet GUI additionally makes the CNN easily accessible to untrained users.

We trained EstrousNet on a random 80% subset of EstrousBank using a ResNet-50-based transfer learning algorithm, yielding test accuracy significantly greater than expert human examiners (**Fig 2**,**3**). Our software incorporates a preloaded trained network for easy adoption, while allowing more advanced users to train their own networks with custom parameters (**Supplementary Fig. S2**). To further improve estrous stage classification, EstrousNet incorporates a cycle fitting algorithm that flags outlier cases in which the deep learning classifications do not line up with an archetypal estrous cycle (**Fig. 4**). In these cases, the GUI gives the user the option to select which classification to accept and incorporates this choice into the net output (**Supplementary Fig. S2**).

Despite our progress in estrous stage classification with EstrousNet and EstrousBank, some limitations remain. Because of the heterogeneity of the training image set, we sacrifice some accuracy for the sake of generalizability. Other CNNs trained on 3 stages from a single dataset therefore exhibit higher validation accuracy in some stages^12,30^. Additionally, the fourth and most transient stage of the estrous cycle, metestrus, yields the lowest test accuracy, as is consistent with previously developed machine learning approaches^12^. Since the presence of both cornified and nucleated epithelial cells in metestrus causes confusion with proestrus, more data will be useful for training CNNs to differentiate between these two stages.

Despite these limitations, misclassifications by EstrousNet remain significantly lower than human experts in diestrus, and similar to expert human coders in proestrus, estrus, and metestrus (**Fig. 2F**). The significantly higher accuracy of diestrus classifications will be useful in flagging the diestrus-proestrus transition, during which estradiol levels spike up to 100-fold ^41,42^. The combination of the easy-to-use software and our highly generalizable algorithm makes EstrousNet an excellent resource for inexperienced classifiers. Our results indicate that human variability remains high even amongst expert coders, highlighting the need for increased inter-lab consistency (**Fig. 2G**). With many experimenters making the transition to using both sexes in rodent studies, generalizable and automated pipelines for tracking estrous stage will be useful for a range of laboratories.

Although 68.3% of EstrousBank images consist of uniform or semi-uniform stains such as crystal violet and H&E, stains designed specifically for hormonal cytodiagnosis offer an opportunity to identify more nuanced biomarkers of the estrous cycle. For instance, Shorr stain makes it possible to distinguish acidophilic and basophilic epithelial cell subtypes, either of which may be more prevalent in the early or late phase of a given estrous stage^40^. Identifying such graded changes in cell type proportionality will be useful for classifying transition stages of the estrous cycle (**Fig. 5**). Characterization of substages will be a step forward in reframing our understanding of the estrous cycle as continuum, instead of a series of discrete stages.

It should be noted that currently there is no ground truth data for cytological stage *in vivo*, as the low concentrations of hormones such as estradiol and progesterone in the bloodstream make daily collection of endocrine data generally intractable in rodents. Although larger rats may have sufficient blood volume for repeated sampling, existing radioimmunoassay techniques are invasive, expensive, and time consuming^43^. At present, most ground truth data from the estrous cycle is derived from terminal experiments in which animals are sacrificed at staggered timepoints and large volumes of blood are used to determine hormone concentration^18,22,41^.

However, advances in biosensors for steroid hormone analysis, including aptamer^44,45^, bioaffinity^46^, and magnetic nanoparticle sensors^47^, offer exciting opportunities for repeated estradiol and progesterone measurements. Additionally, physiological characteristics such as temperature^48^, heart rate^49^, uterine impedance^20^, and blood oxygen content^50^ could be incorporated into estrous stage identification as a proxy for steroid hormone concentrations. As new biomarkers become available, we hope to update EstrousNet to integrate these inputs and further improve the classification accuracy.

Ultimately, it is our goal that accessible technologies for cytological classification will help reduce the exclusion of female animals from scientific studies, a disparity that is especially prevalent in fields such as neuroscience and pharmacology, in which significant sex differences have been described^1,2^. We hope that by continuing to add new cytology images and metadata into our EstrousBank dataset over time, we will be able to bolster our network to identify biological processes that are modulated by steroid hormones.

## METHODS

### Animals

The images in EstrousBank were collected from 5 different labs. Cytology images from the Goard lab were taken from female Thy1-GFP-M transgenic mice and Slc17a7-IRES2-Cre x TITL2-GC6s-ICL-TTA2 double transgenic mice, neither of which showed strain-specific disruptions to the estrous cycle. Animals were housed in cages of up to 5 animals, and singly housed after being surgically implanted with a headplate and cranial window for corresponding imaging experiments. Animals were given food and water ad libitum and kept on a 12 h light/dark cycle. Samples were taken at 16-40 weeks, with a median age of 30 weeks, using vaginal lavage. All animal procedures were approved by the Institutional Animal Care and Use Committee at University of California, Santa Barbara.

Cytology from the Galea Lab was taken from wild-type female Sprague-Dawley rats. Animals were housed in cages of 2-3, given food and water ad libitum, and kept on a 12 h light/dark cycle. Samples were taken at 8-17 weeks of age using vaginal lavage. Older animals were concomitantly involved in behavioral experiments that may have resulted in elevated stress. All experimental procedures were approved by the University of British Columbia Animal Care Committee and were completed in accordance with the Canadian Council on Animal Care guidelines.

Cytology from the Ostroff lab was taken from wild-type female Sprague-Dawley rats. Animals were housed in cages of 2, given food and water ad libitum, and kept on either a 12h or 14:10 light/dark cycle. Cages were filled with autoclaved standard Sani-Chip bedding (Teklad Global, Envigo) and one enrichment device. Samples were taken at 4-14 weeks of age using vaginal swab. All animal protocols were approved by the Institutional Animal Care and Use Committee at the University of Connecticut.

Cytology from the Shansky Lab was taken from wild-type female Long Evans rats. Animals were housed in cages of 2, given food and water ad libitum, and kept on a 12 h light/dark cycle. Samples were taken at average 12-16 weeks using vaginal swab. All animal procedures were approved by the Institutional Animal Care and Use Committee at Northeastern University.

Cytology from the Sutoh lab was taken from wild-type female C57BL/6J mice. Animals were provided food and water ad libitum and kept on a 12 h light/dark cycle. Samples were taken at 5-14 weeks using vaginal swab. All animal-use procedures were in accord with the Guidelines for Animal Experimentation of Showa Pharmaceutical University.

### Vaginal cytology

EstrousBank samples were collected using saline lavage (9.2%) or vaginal swab (90.8%). Vaginal lavage samples were collected using a P200 micropipette. 50 μl sterile saline was pipetted into the vaginal opening and aspirated several times to obtain a sufficient cell count. The sample was pipetted onto a gel subbed microscope slide and allowed to dry 24 h before staining. For vaginal swabs, cotton-tipped swabs were soaked in sterile saline and briefly rolled against the superficial vaginal wall. The epithelial cells on the swab were then transferred to a dry gel subbed glass slide.

Gel subbing was performed in-house using standard IHC protocol to coat glass slides in gelatin/CrK(SO4)2 solution^19^. Staining procedures, including crystal violet, Giemsa, H&E, and Shorr stain, are as described elsewhere^20,40,41,50^.

### EstrousBank curation

The 12,719 images in EstrousBank were contributed from the Goard lab, Ostroff lab, Shansky lab, Galea lab, and Sutoh lab. These labs provided cytology images from a diverse set of histological stains, magnifications, species, and strains (**Supplementary Table S1**). Initial classifications were made based on traditional cell type proportionality, as determined by the source lab. For cross-group consistency, benchmark classifications were made between the experimenters who provided the cytology images and those compiling EstrousBank. Images were classified into a given stage when 2 or more expert coders agreed on a stage classification, including those from transition stages (**Fig. 5**). Images containing excessive debris, n<10 cells, or <300 pixels were excluded (4.6%).

### Image preprocessing

Input images were normalized by aligning maximum peaks of the luminance histograms. Images were then converted to greyscale to allow EstrousNet to generalize onto different stains. After normalization, images in both cohorts were randomly divided into 80% training, 10% validation, and 10% test sets. These images were then split into four quadrants within the same directory. Greyscale images were concatenated into 3D arrays to meet input image size requirements. Images were then stored in an augmented datastore where each image was resized to 224 × 224 × 3.

EstrousNet augmented the quadrupled dataset with X and Y translation, rotation, reflection, and scaling, according to user parameters in EstrousNetTrainNewNet.mlapp, the network training GUI. EstrousNet users can choose to train their own net using custom augmentation parameters in the EstrousNet GUI or load one of our open-source pretrained networks.

### Implementation and training of CNN architectures

The pretrained EstrousNet is based on the ResNet-50 architecture, which yields the highest validation and test accuracy on the EstrousBank images. However, users can choose to train EstrousNet using VGG-19, MobileNet v2, or Inception v3 architectures, the connected layers of which have been prespecified in our code^31–34^. VGG-19 is a network characterized by highly connected convolutional and fully connected layers which enable efficient feature extraction and use Maxpooling for downsampling, unlike the average pooling layers of ResNet50^33^. Compared to ResNet and VGG networks, Inception v3 uses auxiliary classifiers, asymmetric convolutions, and fewer overall parameters for high computational efficiency and low error rates^31^. Finally, MobileNet v2 is a lighter deep neural net ideal that only uses a regular convolution on the first layer of an input image, designed for users with datasets that desire high accuracy with reduced parameters^32^.

In the standard ResNet50 architecture, used here as the base architecture of EstrousNet, nonlinear skip connections and shortcuts are implemented to maintain high performance despite a deep architecture^34^. The residual block on ResNet-50 is defined as follows:

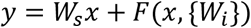

Where *x* is input layer; *y* is output layer; the function *F*(*x*, {*W*_*i*_}) represents the residual mapping to be learned; and *W*_*s*_ is the linear projection performed to match the dimensions of *x* and *F*.

The architecture of ResNet-50 consists of 5 stages, each with a convolution and identity block made up of 3 convolution layers^34,51^. The two initial layers accomplish convolution of size 7 × 7 and max-pooling of size 3 × 3 with a stride of 2^34,51^. Input images are resized to 224×224×3 before undergoing augmentation and training. Training hyperparameters were specified using a Bayesian optimizer, which yielded highest accuracy with an initial learning rate of 1e^-5^ and a mini batch size of 80. Several gradient descent optimization algorithms were tested, including RMSprop, adam, and sgdm, all designed to minimize the loss function of the network. RMSprop exceeded the other algorithms in terms of accuracy when combined with a squared gradient decay of 0.99. Due to the breadth of the input images only 3 epochs were necessary to maintain maximum accuracy, with shuffling occurring every epoch, as well as a piecewise learning rate drop factor of 0.1, the step decay algorithm of which is as follows:

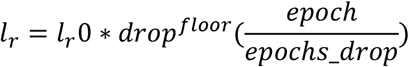

Where *l*_*r*_ is learning rate; *l*_*r*_0 is initial learning rate (here 1e^-5^); drop is the factor by which the learning rate is decreased (here 0.1); *floor* is the minimum learning rate; *epoch* is the current epoch, and *epochs_drop* is the number of epochs after which the step decay will occur (here 1)^52^.

The EstrousNet GUI was developed in MATLAB 2020b (Mathworks, Inc.) using the App Designer platform. EstrousNet was trained using EstrousNetTrainNewNet.mlapp, classification input was given by EstrousNetGUI.mlapp, and classification output was plotted using EstrousNetPlotting.mlapp. The GUI is also used to tune augmentation parameters and number of stages desired for classification.

### Cycle fitting

Here, a custom waveform describing the time course of the estrous cycle was generated using prior publications^10,18,20–23,30,35–38^. The resulting archetypical estrous cycle has a period of 4.87 days (**Fig. 4A**). The stage classifications are ordered diestrus > metestrus in increments of 1.0 starting from 0.5, where 0.0 and 4.0 were defined as the transition stage between metestrus and diestrus (**Fig. 4A**). We fit these points with a two-term polynomial, calculating the coefficients using the temporal midpoints of each stage of the estrous cycle. The periodic waveform is fit to the input data for EstrousNet by shifting the phase by 0.1 cycles and selecting the phase shift with the maximum Pearson’s correlation coefficient (**Fig. 4B**).

Cycle fitting also allowed us to detect anestrous stages (i.e., pseudopregnancy), which are occasionally induced by cytology sampling methods such as vaginal swab and lavage. In our algorithm, the user will receive a pseudopregnancy warning message if the animal has been in diestrus 50% longer than in previous cycles, given that the user specifies sequential data sampling in the GUI (**Fig. 4C-E**). This characterization is consistent with our observation that more than 2 consecutive days of > 90% leukocytes is indicative of an anestrous state (**Fig. 4D**).

### Statistical information

To compare the accuracy of EstrousNet vs trained human examiners, a test set of 400 images was created by randomly selecting 100 images from each of the 4 estrous stages (**Fig. 2D, E)**. Human examiners were expert coders who had each individually classified upwards of 2000 cytology images. EstrousNet was trained on the images in EstrousBank, as described previously, excluding the 400 images in the test set. Benchmark classifications were used as a proxy for ground truth, in the absence of intravenous hormone measurements, as described previously. Accuracy was determined by comparing these ground truth classifications to EstrousNet classifications. These comparisons are represented by a confusion matrix generated in MATLAB (**Fig. 2D,E**).

For statistical analysis, net accuracy and human accuracy vectors for each stage were concatenated and bootstrapped across 5000 iterations to create a normal distribution. Violin plots were made using an open-source MATLAB package^55^. We performed the Fisher’s Exact Test within and across stages to test for significance (**Fig. 2F**).

For out-of-sample testing, three dimensions of sampling were used: stain, species, and subject. For stains and species, each respective category was removed from the training set and set aside for testing. EstrousNet was trained separately for each category on the revised datasets (**Figure 3C-E**). It should be noted that multiple dimensions were nested in our framework, i.e., because each lab group used a different stain for their cytology images, removing any species also removed a set of stains. Accuracy was measured by taking the proportion of EstrousNet classifications that were consistent with benchmark classifications, run across 1000 iterations sampled without replacement to generate standard error. For out-of-sample subject testing 36 individual animals were identified, including 20 WT Sprague Dawley rats and 16 Slc7a7-cre × TITL GCaMP6s B6 mice. k = 6 groups were used for k-fold out of sample cross-validation testing, with 6 subjects in each group. The resulting confusion matrix is an average of the k-fold accuracy results.

ROC curves were generated using the *perfcurve* MATLAB function to generate a logistic regression, then the integral of each curve was taken to calculate the auROC for each stage (**Fig. 3A**). For these curves, true positive was defined as an instance where a given positive stage was correctly classified, whereas false positive was defined as the number of negative stages falsely categorized into a given positive stage.

The sensitivity curve was generated by finding the rate of images in a positive class, i.e., images belonging to a given stage, that were correctly classified as being in that stage (**Fig. 3B**). The specificity curve was generated by finding the rate of images in a negative class, i.e., not part of a given stage, that were correctly classified as not belonging to that stage (**Fig. 3B**). The probability cutoff of 0.26 was defined as the intersection between these two curves (**Fig. 3B**). Pseudopregnancy cell count significance was determined by a two-way ANOVA (**Fig. 4D**).

## DATA AVAILABILITY

All code necessary to run EstrousNet is available at http://github.com/ucsb-goard-lab/EstrousNet. EstrousBank is available in full at [*IDR number to be determined*].

## ACKNOWLEDGEMENTS

We would like to thank Dr. Nina Miolane and Dr. Emily Jacobs for comments on this manuscript, and Dr. Chiro Sutoh for contributing data to the EstrousBank. We would like to thank William Castagna, Marie Karpinska, and Emily Youngblood for assistance collecting cytology samples. This work was supported by the Larry Hillblom foundation (M.J.G.).

## AUTHOR CONTRIBUTIONS

N.S.W. developed EstrousNet; N.S.W and K.K.S. analyzed EstrousNet performance; N.S.W and K.K.S. developed the EstrousNet GUI; G.R., T.H., and N.S.W. classified test images; G.R., T.H., R.M.S, L.A.M.G., L.O, and M.J.G. contributed to EstrousBank curation; N.S.W. and M.J.G. wrote the manuscript; all authors reviewed the manuscript.

## COMPETING INTERESTS

The authors declare no competing interests.

## SUPPLEMENTARY MATERIAL

**Supplementary Figure S1.**
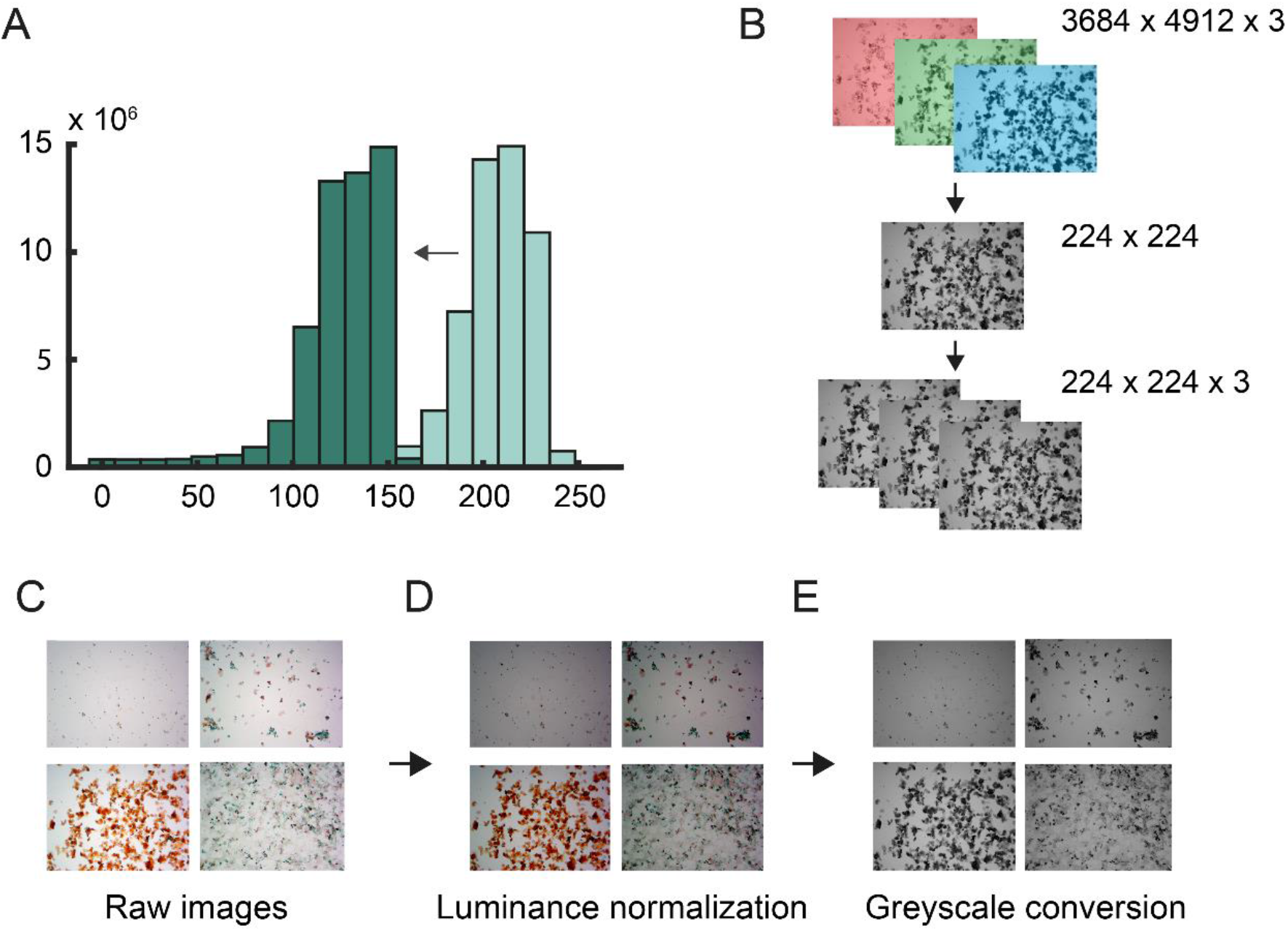
Image preprocessing pipeline. **A**. Intensity histogram of an unprocessed image (light blue), shifted to lower intensity (dark blue) during luminance normalization. **B**. Schematic of image resizing and conversion to grayscale, where 1D grayscale images are concatenated into a 3D array of size 224 × 224 × 3 to match the input requirements of the transfer learning network. **C**. Example unprocessed test images from one estrous cycle. **D**. Raw images with reduced intensity, normalized to the same maximum intensity peak. **E**. Luminance-normalized images converted to 3-channel grayscale.

**Supplementary Figure S2.**
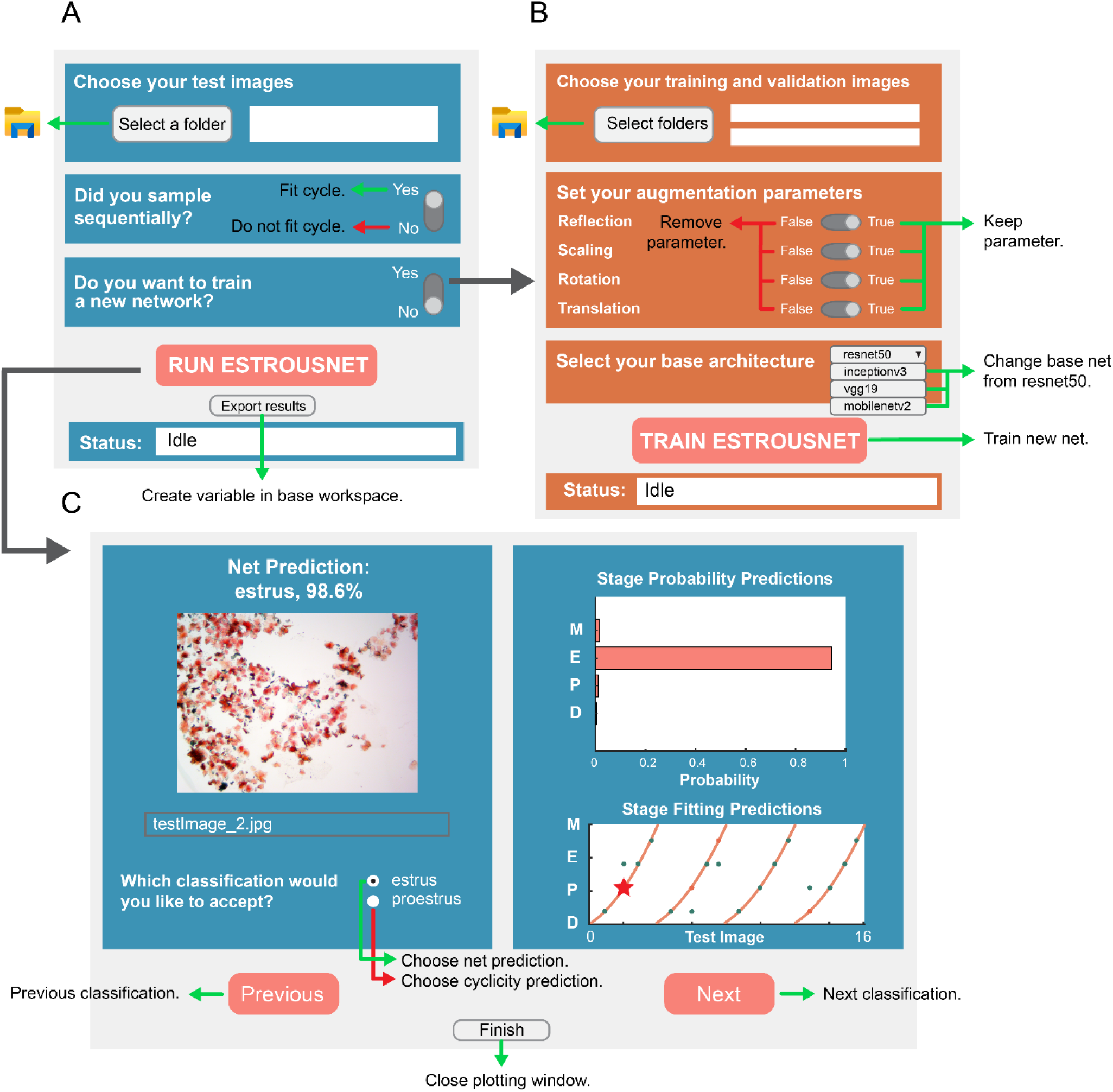
Illustration of the EstrousNet user interface (GUI). **A**. The EstrousNet classification GUI: the user selects a folder of test images which are automatically classified and plotted. The user also selects whether images were sampled sequentially, which will determine whether net classifications are fit to an archetypical cycle. **B**. The EstrousNet training GUI: if the user selects that they would like to train a new network, it will launch the training GUI. This GUI lets the user select folders with training and validation images, as well as custom augmentation parameters, and once training is finished will save the trained network and training data to the current directory. **C**. The EstrousNet plotting GUI: once the classification GUI is used to select test images, the plotting GUI will display the results of the net classifications. If images were taken in sequence, the plotting GUI will fit the images to an archetypal cycle, and for any images where the cyclicity and net classifications disagree, the user can choose to manually select the preferred classification.

**Supplementary Table S1.** Summary of EstrousBank images from multiple labs. Metrics for the images included in the open-source image repository EstrousBank, subdivided by the groups contributing the raw images.

| Source Lab | Magnification | Stain | Species | Strain | # images | % of total images |
| --- | --- | --- | --- | --- | --- | --- |
| Galea | 20X | Cresyl Violet | Rat | Sprague Dawley WT | 145 | 1.14 |
| Goard | 10X | H&E, Shorr Stain | Mouse | Thy1-GFP-M (Jax Stock #007788), Slc7a7-cre (Jax Stock #023527) x TITL-GCaMP6s (Jax Stock #024104) C57BL/6J | 1024 | 8.05 |
| Ostroff | 10X | H&E, Shorr Stain | Rat | Sprague Dawley WT | 6277 | 49.35 |
| Shansky | 10X | Crystal Violet | Rat | Long Evans WT | 1954 | 15.36 |
| Sutoh | 10X | Giemsa | Mouse | C57BL/6J WT | 3319 | 26.09 |

## Notes

### Competing Interest Statement

The authors have declared no competing interest.

## REFERENCES

1. Woitowich, N. C., Beery, A. K. & Woodruff, T. K. A 10-year follow-up study of sex inclusion in the biological sciences. eLife 9, 1–8 (2020).

2. Shansky, R. M. & Murphy, A. Z. Considering sex as a biological variable will require a global shift in science culture. Nature Neuroscience 2021 24:4 24, 457–464 (2021).

3. Pritschet, L. et al. Functional reorganization of brain networks across the human menstrual cycle. NeuroImage 220, 117091 (2020).

4. Woolley, C. S. & McEwen, B. S. Roles of estradiol and progesterone in regulation of hippocampal dendritic spine density during the estrous cycle in the rat. Journal of Comparative Neurology 336, (1993).

5. Woolley, C. S. & McEwen, B. S. Estradiol mediates fluctuation in hippocampal synapse density during the estrous cycle in the adult rat. Journal of Neuroscience 12, (1992).

6. Kim, J. & Frick, K. M. Distinct effects of estrogen receptor antagonism on object recognition and spatial memory consolidation in ovariectomized mice. Psychoneuroendocrinology 85, (2017).

7. Galea, L. A. M., Perrot-Sinal, T. S., Kavaliers, M. & Ossenkopp, K. P. Relations of hippocampal volume and dentate gyrus width to gonadal hormone levels in male and female meadow voles. Brain Research 821, (1999).

8. Hara, Y., Waters, E. M., McEwen, B. S. & Morrison, J. H. Estrogen Effects on Cognitive and Synaptic Health Over the Lifecourse. Physiological Reviews 95, 785 (2015).

9. Frick, K. M., Kim, J., Tuscher, J. J. & Fortress, A. M. Sex steroid hormones matter for learning and memory: estrogenic regulation of hippocampal function in male and female rodents. Learning & Memory 22, 472–493 (2015).

10. Byers, S. L., Wiles, M. v., Dunn, S. L. & Taft, R. A. Mouse Estrous Cycle Identification Tool and Images. PLOS ONE 7, e35538 (2012).

11. Long, J. A. & Evans, H. M. The oestrous cycle in the rat and its associated phenomena. (University of California Press, 1922).

12. Sano, K. et al. Deep learning-based classification of the mouse estrous cycle stages. Scientific Reports 2020 10:1 10, 1–8 (2020).

13. Iqbal, J. et al. Estradiol Alters Hippocampal Gene Expression during the Estrous Cycle. Endocrine Research 45, 84–101 (2020).

14. Vastagh, C. & Liposits, Z. Impact of proestrus on gene expression in the medial preoptic area of mice. Frontiers in Cellular Neuroscience 11, 183 (2017).

15. Woolley, C. S., Gould, E., Frankfurt, M. & McEwen, B. S. Naturally occurring fluctuation in dendritic spine density on adult hippocampal pyramidal neurons. Journal of Neuroscience 10, 4035–4039 (1990).

16. Kashuba, A. D. M. & Nafziger, A. N. Physiological Changes During the Menstrual Cycle and Their Effects on the Pharmacokinetics and Pharmacodynamics of Drugs. Clinical Pharmacokinetics 1998 34:3 34, 203–218 (2012).

17. Gong, S. et al. Dynamics and Correlation of Serum Cortisol and Corticosterone under Different Physiological or Stressful Conditions in Mice. PLOS ONE 10, e0117503 (2015).

18. Haim, S., Shakhar, G., Rossene, E., Taylor, A. N. & Ben-Eliyahu, S. Serum levels of sex hormones and corticosterone throughout 4-and 5-day estrous cycles in Fischer 344 rats and their simulation in ovariectomized females. Journal of Endocrinological Investigation 2003 26:10 26, 1013–1022 (2014).

19. Westwood, F. R. The Female Rat Reproductive Cycle: A Practical Histological Guide to Staging. Toxicologic Pathology 36, 375–384 (2008).

20. Ajayi, A. F. & Akhigbe, R. E. Staging of the estrous cycle and induction of estrus in experimental rodents: an update. Fertility Research and Practice 2020 6:1 6, 1–15 (2020).

21. Cora, M. C., Kooistra, L. & Travlos, G. Vaginal Cytology of the Laboratory Rat and Mouse: Review and Criteria for the Staging of the Estrous Cycle Using Stained Vaginal Smears. Toxicologic Pathology 43, 776–793 (2015).

22. Goldman, J. M., Murr, A. S. & Cooper, R. L. The rodent estrous cycle: characterization of vaginal cytology and its utility in toxicological studies. Birth Defects Research Part B: Developmental and Reproductive Toxicology 80, 84–97 (2007).

23. Paccola, C. C. et al. The rat estrous cycle revisited: a quantitative and qualitative analysis. Animal Reproduction 10, 677–683 (2018).

24. de Fauw, J. et al. Clinically applicable deep learning for diagnosis and referral in retinal disease. Nature Medicine 2018 24:9 24, 1342–1350 (2018).

25. Esteva, A. et al. Dermatologist-level classification of skin cancer with deep neural networks. Nature 2017 542:7639 542, 115–118 (2017).

26. Gurovich, Y. et al. Identifying facial phenotypes of genetic disorders using deep learning. Nature Medicine 2019 25:1 25, 60–64 (2019).

27. Shen, D., Wu, G. & Suk, H. il. Deep Learning in Medical Image Analysis. Annual Review of Biomedical Engineering 19, 221–248 (2017).

28. Hu, J. et al. Iterative transfer learning with neural network for clustering and cell type classification in single-cell RNA-seq analysis. Nature Machine Intelligence 2020 2:10 2, 607–618 (2020).

29. Yao, K., Rochman, N. D. & Sun, S. X. Cell Type Classification and Unsupervised Morphological Phenotyping From Low-Resolution Images Using Deep Learning. Scientific Reports 2019 9:1 9, 1–13 (2019).

30. Pantier, L., Li, J. & Christian, C. Estrous Cycle Monitoring in Mice with Rapid Data Visualization and Analysis. Bio-Protocol 9, (2019).

31. Szegedy, C., Vanhoucke, V., Ioffe, S., Shlens, J. & Wojna, Z. Rethinking the Inception Architecture for Computer Vision. Proceedings of the IEEE Computer Society Conference on Computer Vision and Pattern Recognition. 2818–2826 (2015).

32. Sandler, M., Howard, A., Zhu, M., Zhmoginov, A. & Chen, L. C. MobileNetV2: Inverted Residuals and Linear Bottlenecks. Proceedings of the IEEE Computer Society Conference on Computer Vision and Pattern Recognition 4510–4520 (2018).

33. Simonyan, K. & Zisserman, A. Very Deep Convolutional Networks for Large-Scale Image Recognition. 3rd International Conference on Learning Representations, ICLR 2015 - Conference Track Proceedings (2014).

34. He, K., Zhang, X., Ren, S. & Sun, J. Deep Residual Learning for Image Recognition. Proceedings of the IEEE Computer Society Conference on Computer Vision and Pattern Recognition, 770–778 (2015).

35. Yoshinaka, K. et al. Effect of different light–dark schedules on estrous cycle in mice, and implications for mitigating the adverse impact of night work. Genes to Cells 22, 876–884 (2017).

36. van Goethem, N. P. et al. Object recognition testing: Rodent species, strains, housing conditions, and estrous cycle. Behavioural Brain Research 232, 323–334 (2012).

37. Caligioni, C. S. Assessing Reproductive Status/Stages in Mice. Current Protocols in Neuroscience 48, A.4I.1-A.4I.8 (2009).

38. Spencer, J. L., Waters, E. M., Milner, T. A. & McEwen, B. S. Estrous cycle regulates activation of hippocampal Akt, LIM kinase, and neurotrophin receptors in C57BL/6 mice. Neuroscience 155, 1106–1119 (2008).

39. Kiyonari, H. et al. Targeted gene disruption in a marsupial, Monodelphis domestica, by CRISPR/Cas9 genome editing. Current Biology 31, 3956-3963.e4 (2021).

40. Shorr, E. A New Technic for Staining Vaginal Smears: III, a Single Differential Stain. Science 94, 545–546 (1941).

41. McLean, A. C., Valenzuela, N., Fai, S. & Bennett, S. A. L. Performing Vaginal Lavage, Crystal Violet Staining, and Vaginal Cytological Evaluation for Mouse Estrous Cycle Staging Identification. JoVE (Journal of Visualized Experiments) e4389 (2012).

42. Singletary, S. J. et al. Lack of Correlation of Vaginal Impedance Measurements with Hormone Levels in the Rat. Contemporary topics in laboratory animal science / American Association for Laboratory Animal Science 44, 37 (2005).

43. Skenandore, C. S., Pineda, A., Bahr, J. M., Newell-Fugate, A. E. & Cardoso, F. C. Evaluation of a commercially available radioimmunoassay and enzyme immunoassay for the analysis of progesterone and estradiol and the comparison of two extraction efficiency methods. Domestic Animal Endocrinology 60, 61–66 (2017).

44. Contreras Jiménez, G. et al. Aptamer-based label-free impedimetric biosensor for detection of progesterone. Analytical Chemistry 87, (2015).

45. Nameghi, M. A. et al. An ultrasensitive electrochemical sensor for 17β-estradiol using split aptamers. Analytica Chimica Acta 1065, (2019).

46. De, S., Macara, I. G. & Lannigan, D. A. Novel biosensors for the detection of estrogen receptor ligands. Journal of Steroid Biochemistry and Molecular Biology 96, (2005).

47. Jia, Y. et al. Magnetic nanoparticle enhanced surface plasmon resonance sensor for estradiol analysis. Sensors and Actuators B: Chemical 254, 629–635 (2018).

48. Kent, S., Hurd, M. & Satinoff, E. Interactions between body temperature and wheel running over the estrous cycle in rats. Physiology & Behavior 49, 1079–1084 (1991).

49. Takezawa, H., Hayashi, H., Sano, H., Saito, H. & Ebihara, S. Circadian and estrous cycle-dependent variations in blood pressure and heart rate in female rats. 267, (1994).

50. Mitchell, J. A. Y. J. Intrauterine oxygen tension during the estrous cycle in the rat: its relation to uterine respiration and vascular activity. Endocrinology 83, 701–705 (1968).

51. Gronroos, M. & Kauppila, O. Hormonal-cyclic changes in rats under normal conditions and under stress as revealed by vaginal smears after Shorr staining. Acta endocrinologica 32, (1959).

52. Ge, R., Kakade, S. M., Kidambi, R. & Netrapalli, P. The Step Decay Schedule: A Near Optimal, Geometrically Decaying Learning Rate Procedure for Least Squares. Advances in Neural Information Processing Systems 32, (2019).

